# Metagenome-based microbial community analysis of urine-derived fertilizer

**DOI:** 10.1101/2023.12.05.570237

**Authors:** Nebiyat N. Woldeyohannis, Adey F. Desta

## Abstract

The present study aimed to understand the bacterial portion of the microbial community composition and dynamics of plasmid-mediated antimicrobial resistant genes during the optimized process of struvite production from composite human urine. Samples for DNA extraction was collected from fresh urine, stored urine and struvite during the process of struvite production. Shotgun metagenomic analysis was employed to understand the bacterial community. The most dominant phyla in the fresh and stored urine samples were Pseudomonadata, which comprised of 60% and 43% respectively, followed by Bacillota, comprised of 25% and 39% respectively. The struvite sample was dominated by the phylum Bacilliota (61%), Pseudomonadota (18%) and bacteroidota (12%). The members of the above phyla persisted in dominating each sample accordingly. Member of the family Morganellaceae was dominant in the fresh sample while the stored urine and struvite samples were dominated by the family Clostridiaceae. A decrease of members of the class Gammaproteobacteria was observed from the fresh to the struvite sample though not statistically significant. The genus *Pseudomonas* remained to be the most dominant member of Gammaproteobacteria in the fresh and stored urine sample with OTU count of 12,116 and 6,155 with a marked decrease by half in the stored sample. On the other hand, members of the genera *Clostridium, Enterococcus, Bacteroides* in the stored samples and *Clostridium, Alkaliphilus* and *Pseudomonas* in the struvite samples were dominant. Ninety-six percent of the identified genera were shared in all the samples and the antimicrobial resistance genes (ARGs) identified in the fresh urine were shared by the struvite but not by the stored urine (eg. *sul, cat, aph* and *aac* members). the presence of high abundance of ARGs in struvite needs attention in the persistence and transmissibility of the ARGs before application for agriculture.

## Introduction

Urine contains nutrients essential for plant growth and development. Despite the idea that urine in the human body is sterile, studies have evidenced that urine hosts a set of microbes from the anatomical reservoir to its release out (Hilt et al., 2014). Depending on the health status of the individual, a variety of microbes have been reported in human urine (Xu et al., 2022; Cao *et al*., 2022; Andreu, 2005). Urine bacterial composition differs by gender, age (Lewis et al., 2013), contamination with fecal matter when voided (Bischel et al., 2015; Ahmed et al., 2017). Above all, these differences are better captured via applying nucleic acid-based methods (Jung et al., 2019). With the emergence of culture-independent approaches and the subsequent human microbiome project (Turnbaugh et al., 2007), studies revealed the presence of a unique microbiota in the urinary tract system (reviewed in Zuber et al., 2023), the microbial structure and composition is also reported to be subject to change with age, sex and health status (Halinski et al., 2023, Pearce et al., 2014). Next-Generation Sequencing (NGS) targeting 16S rRNA was the most widely used approach to determine the bacterial portion of the urinary microbiome (Perez-Carrasco et al., 2021, Siddiqui et al., 2011, Lahr et al., 2016). The few studies that involved shotgun metagenomic approach revealed the microbial, archaeal, viral and eukaryote members of the urinary microbiome in addition to the microbial profiles in clinically asymptomatic individuals and symptomatic patients (Hasman et al., 2014 and Moustafa et al., 2018). The involvement of unculturable microbes in specific urogenital disease conditions was revealed by various researchers (Ackerman and Chai, 2019; Fricke et al., 2014; Siddiqui et al., 2012).

Besides the clinical significance, metagenomic analysis of voided urine is important to get a snapshot of the microbial community in urine-derived fertilizer and evaluate the safety of the product (Zhou et al., 2019; Sun et al 2021; Marti et al., 2013; Zhang et al., 2019 and Black et al., 2021). Bishel, et al. (2016), reported flowcytometry and culture-based approaches on the survival of bacteria inoculated for tracking. On the other hand, 16S RNA amplicon sequencing was performed on fresh and stored urine (Lahr et al., 2016).

Metagenomic approach provides an overarching view of the microbiota of urine as it goes all the way to struvite production. In previous work, we examined the resistome carried by mobile genetic elements of bacterial or viral origins (Woldeyohannis and Desta, 2023). This study included the bacterial community and diversity analysis that may show insights into public health and community health risk. Addressing the issue of efficient urine sanitization is important if it is to be applied to improve productivity with minimal risk to human and the ecosystem (Zhou et al., 2019; Sun et al 2021; Black et al., 2021).

Thus, this study is intended to show the dynamics of microbial population through struvite formation. We hypothesized that storage of urine before the process of struvite production will have an impact on the quality of the struvite in terms of the microbial content. Storage is known to reduce enteric microbes in urine. Besides, struvite drying was also presumed to have an impact on reduction of microbes as a result of the reduced water activity. The objective of this study is to show the microbial structure and dynamics from fresh urine to struvite step.

## Materials and Methods

### Urine sample collection process and struvite production

Sixty liter of urine was collected from voluntary passersby in one of the busy roads of Addis Ababa, piazza, using disinfected jerricans in October 2018. The filed jerrycans were automatically transported to the bio-instrumentation laboratory of Addis Ababa University at ambient temperature. For this study, urine samples were considered “fresh” within a maximum of one hour from the time of onsite urine collection from which 2L was taken and stored at −20°C until DNA extraction. Stored urine sample was collected from the urine that was tight sealed and stored at room temperature in the laboratory for 20 days. Struvite sample was collected from the struvite produced following the optimized protocol developed previously (Woldeyohannis and Desta, 2023). All the samples were kept in −20°C until the subsequent steps.

### DNA extraction and sequencing

Bacterial genomic DNA was extracted from the fresh, stored urine and struvite. The liquid urine urine samples were thawed at 8°C and then were shaken before use. Five milliliters of each of the urine samples were poured into the 15 ml falcon tubes. Samples of urine were kept on ice and 2ml cold extraction buffer (0.15M NaCl, 0.1M Na 2 EDTA, pH-8) was added. Then, samples were ultrasonicated (Bandelin Sonopuls, Germany) on ice at power 20KHz, 45sec every 15min interval (Geissler et al., 2012). To the sample on ice, 2ml of lysis buffer (0.1M NaCl, 0.5M Tris, 10% SDS pH-8, where it is adjusted by HCl) was added (Tsai and Olson, 1991). Then, 40μl of proteinase K (20mg/ml) was added into each mixture and incubated at 60°C for 1hr (Silva et al., 2013). Equal volume of phenol:chloroform:isoamyl alcohol (24:25:1) was added and the samples were centrifuged at 5000 rpm for 10 min. After transferring the aqueous phase into a new tube, 2.5ml of chloroform was added and centrifuged as previously done. To the collected aqueous phase, 2ml cold isopropanol (−20°C) was mixed for washing at 5000rpm for 10 min. Supernatant was removed and pellet was washed by 2ml cold 70% ETOH. Dried pellets were then dissolved with sterile 20μl Mili-Q water (Applichem, Germany). The extracted DNA was transferred into eppenddorff tubes and stored at −20°C after DNA quality and quantity was measured. For the extraction of the struvite sample, NucleoSpin Soil kit (Macherey-Nagel) was used on 2.5 g sample following the kit instruction, quality check and stored at −20°C until further analysis.

Sequencing was done using Illumina HiSeq X with sequencing shotgun library preparation using the KAPA HyperPrep Kit (Kapa Biosystems) following the manufacturer’s instruction. Adapter ligation, amplification was done with SPRIselect Reagent (Beckman Coulter, CA, USA). Samples were pooled and sequenced for 150 bp read length in paired-end mode, with an intended output of 20 million reads per sample.

## Data Availability

Paired reads from the three samples were deposited in the NCBI with the accession ID SRX19444964 to SRX19444967 and Biosamples assigned from SAMN33336460 to SAMN33336463.

## Sequence analysis

Read quality was checked by the FastQC (V 0.11.7) tool using Phred score of 30. Adapter trimming was done using trimmomatic (V 0.33) and assembly using velvet (V 1.2.9) using values N50, q25. For the community analysis, targeted BLAST (V 2.12.0) tool was used. Annotation and OTU assignment was done using MGRAST, then cleaned-up using openrefine (V 3.2). OTU and taxonomy were analyzed using phyloseq in R (V 4.2.2). Additionally, packages such as ggplot2, tibble and dplyr were applied for data visualization.

## Results

### Phylum-level microbial composition

A total of about 201,132, 316,150 and 246,769 OTUs were detected from fresh urine, stored urine and struvite respectively. The most dominant bacterial phylum in fresh and stored urine was Pseudomonadota (synonym Proteobacteria) with 120,925 OTUs (60 % of the total OTU within sample) in fresh urine and 143,409 OTUs (45 %) in stored urine (Fig. 1). In the struvite sample, Bacillota was the most dominant phylum with 150,200 OTUs, constituting 60% of the total OTU in the struvite sample. Bacillota was also the second most abundant phyla both in fresh and stored urine samples, comprising 52,090 (26%) and 124,273 (39%) OTUs respectively (Fig. 1). The second most abundant phylum in the struvite sample was Pseudomonadota, with 150,200 OTUs that constituted 18% of the total identified OTU.

**Figure 1.**
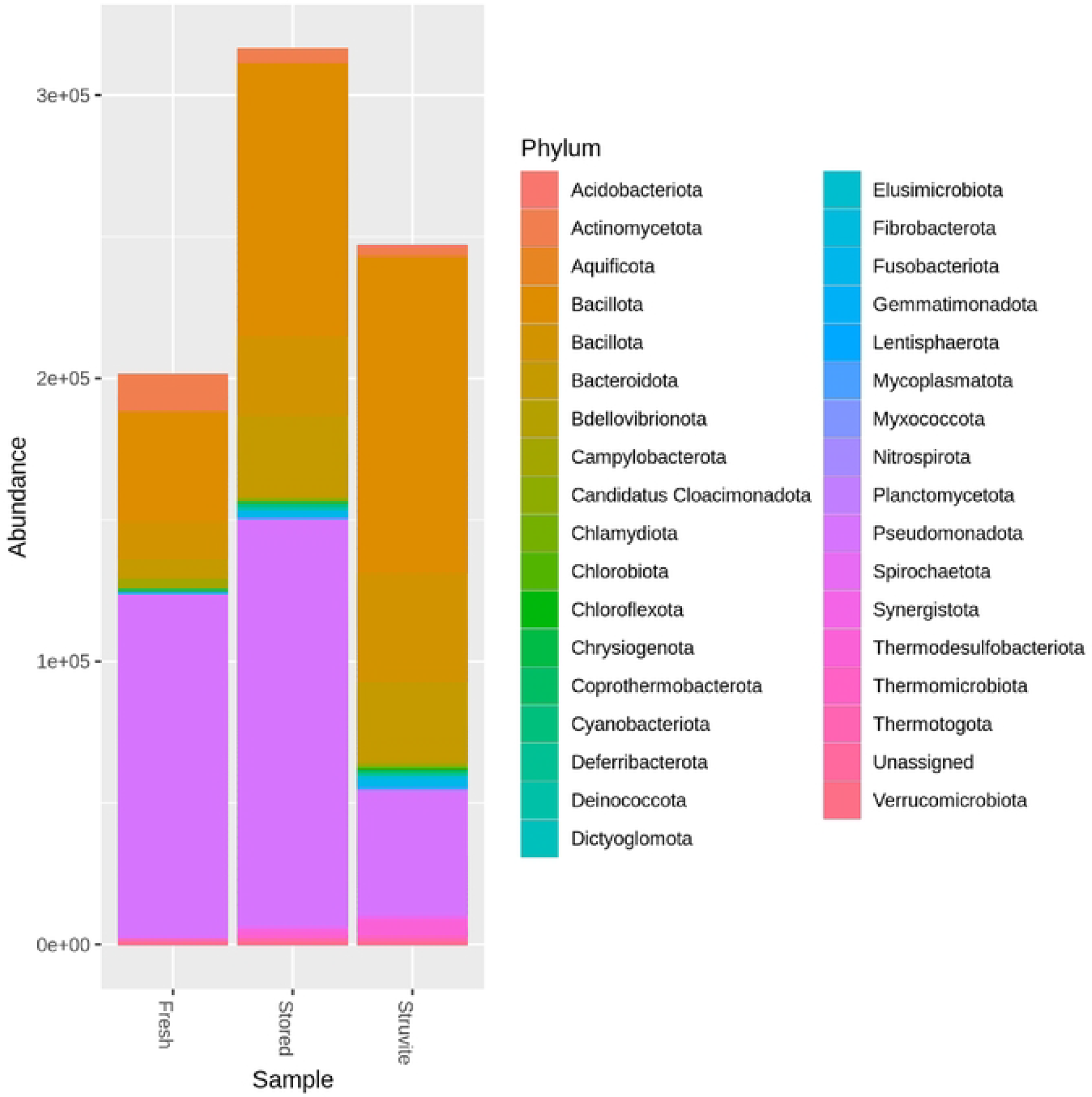
Distribution of bacterial phyla in fresh urine, stored urine and struvite samples

Figure 1. Distribution of bacterial phyla found in fresh urine, stored urine and struvite

In the process of struvite production from fresh urine, the dominant phylum Pseudomonadota showed a decreasing trend in the struvite sample after a slight increase in the stored urine sample (Fig. 2). On the other hand, the phylum Bacillota doubled in abundance in the stored urine samples which then increased by nearly one-fifth in the struvite sample. Likewise, the phylum Bacteroidota showed an increasing trend. In all the samples, the increase and decrease were not statistically significant (p>0.05) based on Mann-Witney test.

**Figure 2.**
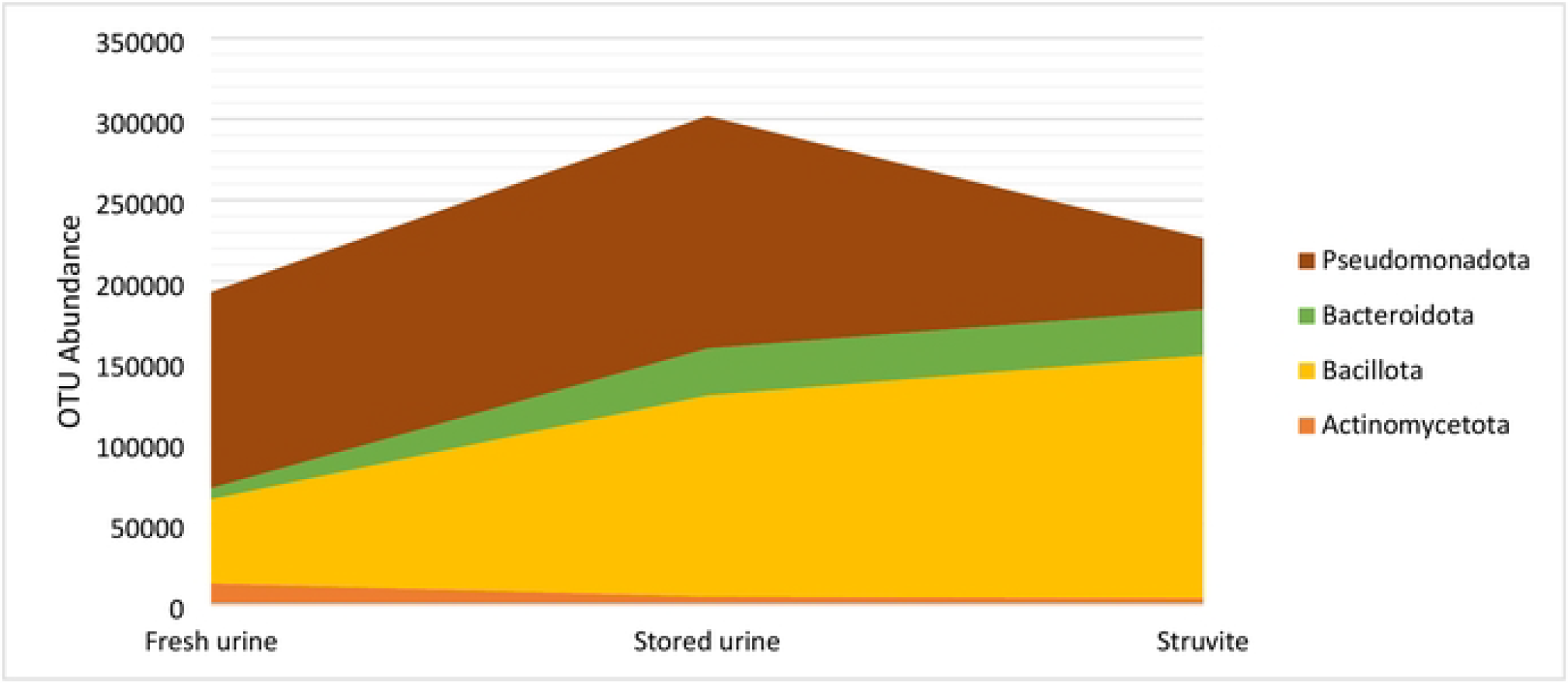
Abundance of the dominant phyla across samples in the process of struvite production

### Class-level microbial distribution in fresh, stored urine and struvite samples

In the fresh urine sample, the class Alphaproteobacteria was the most dominant class with OTU count of 60,773 which increased to 98,064 in stored urine while its OTU count was only 2,823 in struvite. Rather in the struvite sample, the most abundant class of bacteria was Clostridia with 92, 166 OTU count. Clostridia in fresh and stored urine samples decreased to 9,794 and 46,352 OTUs respectively (Figure 3). The second most abundant class in the fresh urine sample was Gammaproteobacteria with 52,815 OTU count, which decreased to 23,804 and 23,495 OTU count in stored urine and struvite respectively. Bacilli was the third most abundant class in the fresh urine sample with OTU count of 41,025. This class was found to increase in stored urine in to 63,443 OTU but in struvite with a slight decrease of 42,405 OTU count (Figure 3).

**Figure 3.**
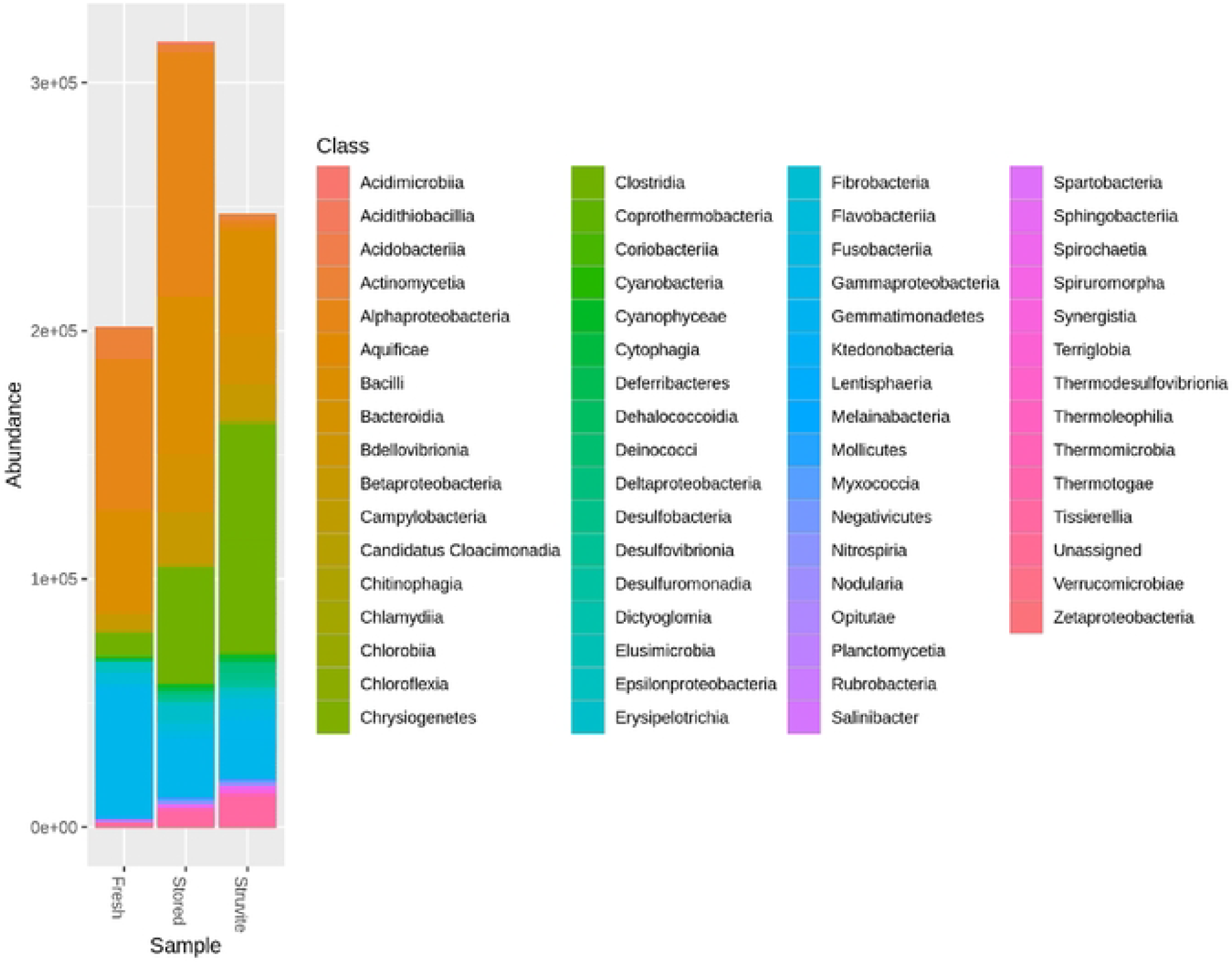
Bacterial classes identified in fresh, stored urine and struvite samples

### Family-level microbial distribution in fresh, stored urine and struvite samples

In the fresh urine sample, the family Morganellaceae was the most abundant family constituting 16,304 OTU count, followed by Pseudomonadaceae (12,647 OTUs), Caulobacteriaceae (12,450 OTUs) and Enterococcaceae (10,738 OTUs).

In the stored urine sample, the most dominant OTU belonged to the family Clostridiaceae, with 27,764 OTU counts constituting 9 % of the total OTU, followed by Sphingomonadaceae with 18,688 OTU counts and Enterococcaceae with 16,475 OTU counts. The members Moraxellaceae 1,981 OTUs), Xanthomonadaceae (1,595 OTUs) and Enterobactereaceae (1,556 OTUs) constituted 7 % each in the stored urine samples. Pasteurellaceae (1,137 OTUs) and Morganellaceae (1,132 OTUs) constituted 5 % each. Vibrionaceae (1,044 OTUs), Aeromonadaceae (926 OTUs), Shewanellaceae (877 OTUs), Ectothiorhodspiraceae (845 OTUs) constituted 4% each. Yersiniaceae and Altreromonadaceae were observed to be 3 % each. Legionellaceae and Psychromonadaceae were observed to be 1 % each.

Similar to the stored urine sample the struvite sample was dominated by the family Clostridiaceae, constituting 59,809 OTU counts with 24% abundance followed by Peptoniphilaceae (15,204 OTUs, 6%) and Bacillaceae (13,276 OTUs, 5%) (Supplementary table 1). the family Pseudomonadaceae (11,716 OTUs) and Bacteroideceae (10,387 OTUs) constituted nearly 5% of the total bacteria in the struvite sample while Alcaligenaceae, Thermoanaerobacteraceae, Enterococcaceae and Lactobacillaceae constituted nearly 3% each (Fig. 4).

**Figure 4.**
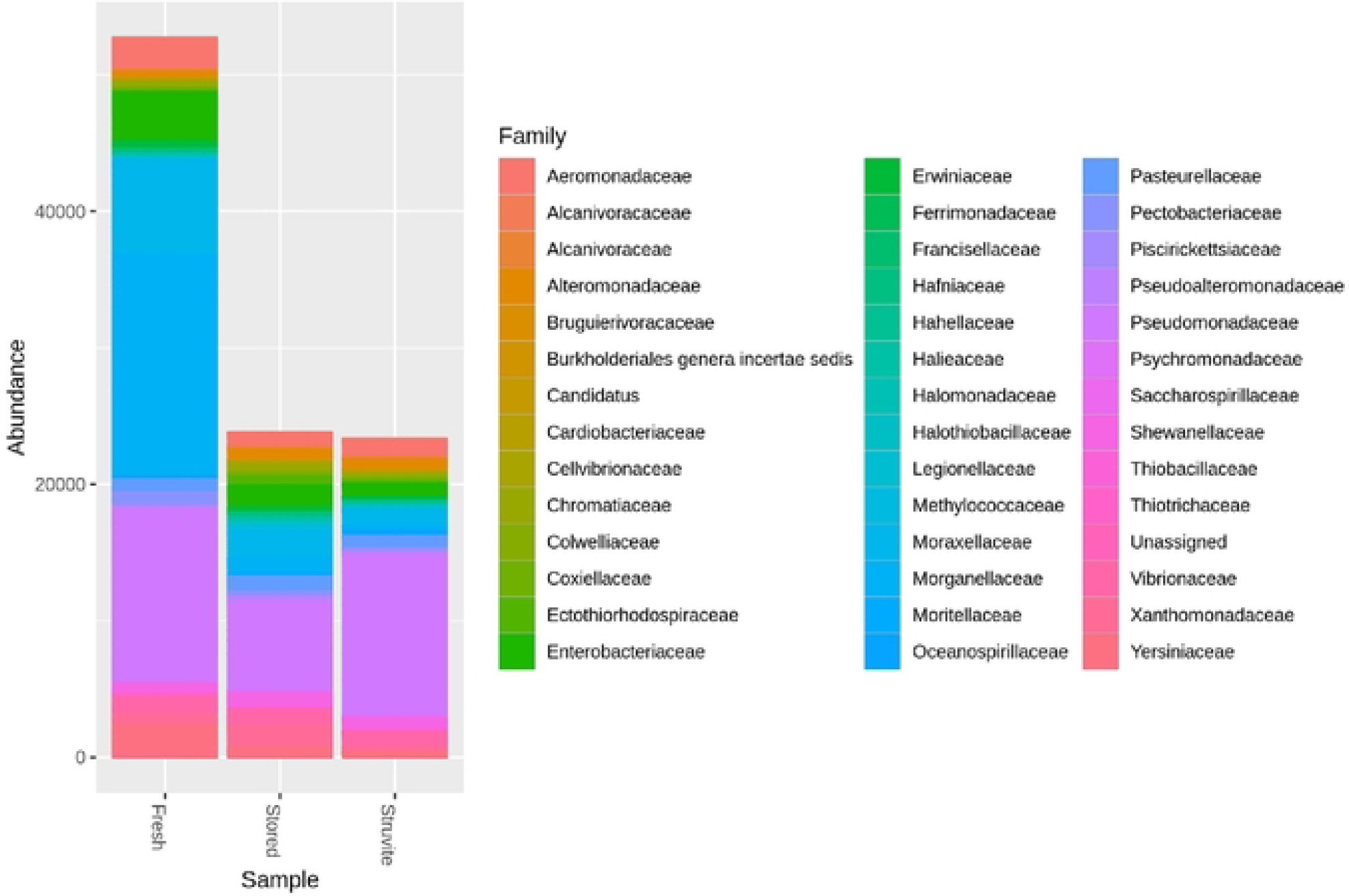
Abundance of members of Gamma proteobacteria in fresh, stored urine and struvite samples

A total of 201,129 OTU counts of families belonging to Gammaproteobacteria were found. In the fresh urine sample, 52,730 OTU count was identified which decreased to 23,800 and 23,339 OTU counts in the stored urine and struvite samples respectively. In the fresh urine sample, the family Yersiniaceae (2375 OTUs) and Aeromonadaceae (2,297 OTUs) constituted nearly 5% of the total OTUs while each of the Vibrionaceae (1,164 OTUs), Pasteurellaceae (950 OTUs), Shewanellaceae (793 OTUs) constituted 2 % of the total OTU counts (Fig. 4). Erwiniaceae, Chromatiaceae, Alteromonadaceae, Ectothorhodospiraceae constituted 1 % each, among the others.

### Genus-level microbial distribution with emphasis on members of the class Gammaproteobacteria

The genus-level OTU of class Gammaproteobacteria decreased by 54% from 52,730 in fresh urine to 23,800 in stored urine and in struvite to 23,339 OTU counts. In the fresh urine, the genus *Pseudomonas* was dominant with OTU count of 12,116, followed by *Providencia* (8,653 OTUs) and *Psychrobacter* (3,646 OTUs) (Fig. 5). In all the identified bacterial OTU, members of the genus *Brevundimonas* constituted 8,273 OTU count, followed by the genera *Rhodopseudomonas* and *Streptococcus*, constituting nearly 4% of the total genera within the fresh sample. The relative abundance of members of the genera Bacillus, *Clostridium, Proteus, Caulobacter, Lactobacillus, Listeria* and *Escherichia* was less than 3% in the fresh urine sample (Supplementary table 2).

**Figure 5.**
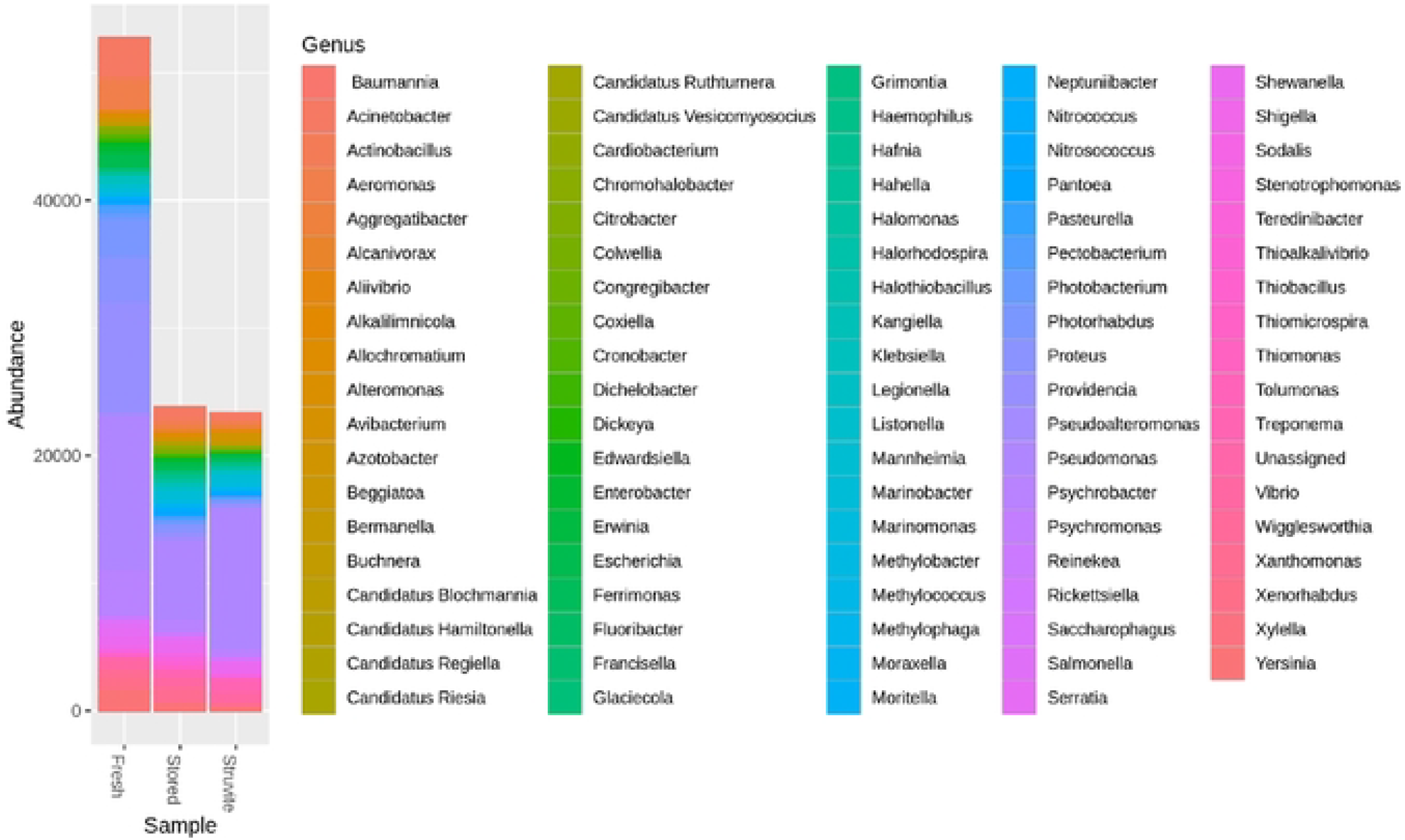
Abundance of genera belonging to the class Gammaproteobacteria in fresh, stored urine and struvite samples

In the stored urine sample, the genus *Clostridium* constituted 18,577 OTU counts making it the most dominant genus followed by *Enterococcus* (15,052 OTUs) and *Bacteroides* (10,288 OTUs). Members of the genera with relative abundance of nearly 2% included *Bacillus, Rhodopseudomonas, Sphingomonas, PseuIdomonas, StreptococcIus* and *Lactobacillus* in decreasing order. The genus *Pseudomonas* remained to be the most dominant member of Gammaproteobacteria in the stored urine sample with OTU count of 6,155 which showed a decrease by nearly 50% from the fresh urine sample (OTU 12,116) and increased by 43% to 10,955 in the struvite sample (Fig. 5).

The most abundant genus in the struvite sample was *Clostridium*, with 36,456 OTU counts. The second and third most dominant genera in struvite were *Alkaliphilus* and *Pseudomonas*, with 16,172 and 10,840 OTU count respectively. The genera *Bacillus, Bacteroides, Enterococcus, Bordetella, Streptococcus, Parabacteroides* and *Lactobacillus* constituted nearly 2% of the total abundance within the sample. Among the members of Gammaproteobacteria, the genera *Treponema* and *Shewanella* were the second and third most abundant genera in the struvite sample (Fig. 5).

In our study, the majority of the genera (96%) were shared among the three samples with a very small proportion of unique genera. It was also observed that Fresh urine had no unique genera while stored urine and struvite samples harbor six unique genera each (Fig. 6). The number of genera shared by any two of the samples was observed to be not more than four. In the stored urine sample, the genera *Leptospirillum, Borreliella, Brugia, Orientia, Prausnitzii* and *Fluoribacter* were the unique ones while in the struvite sample, the genera *Borrelia, Collimonas, Hafinia, Marinococcus, Ornithobacterium* and *Pediococcus*.

**Figure 6.**
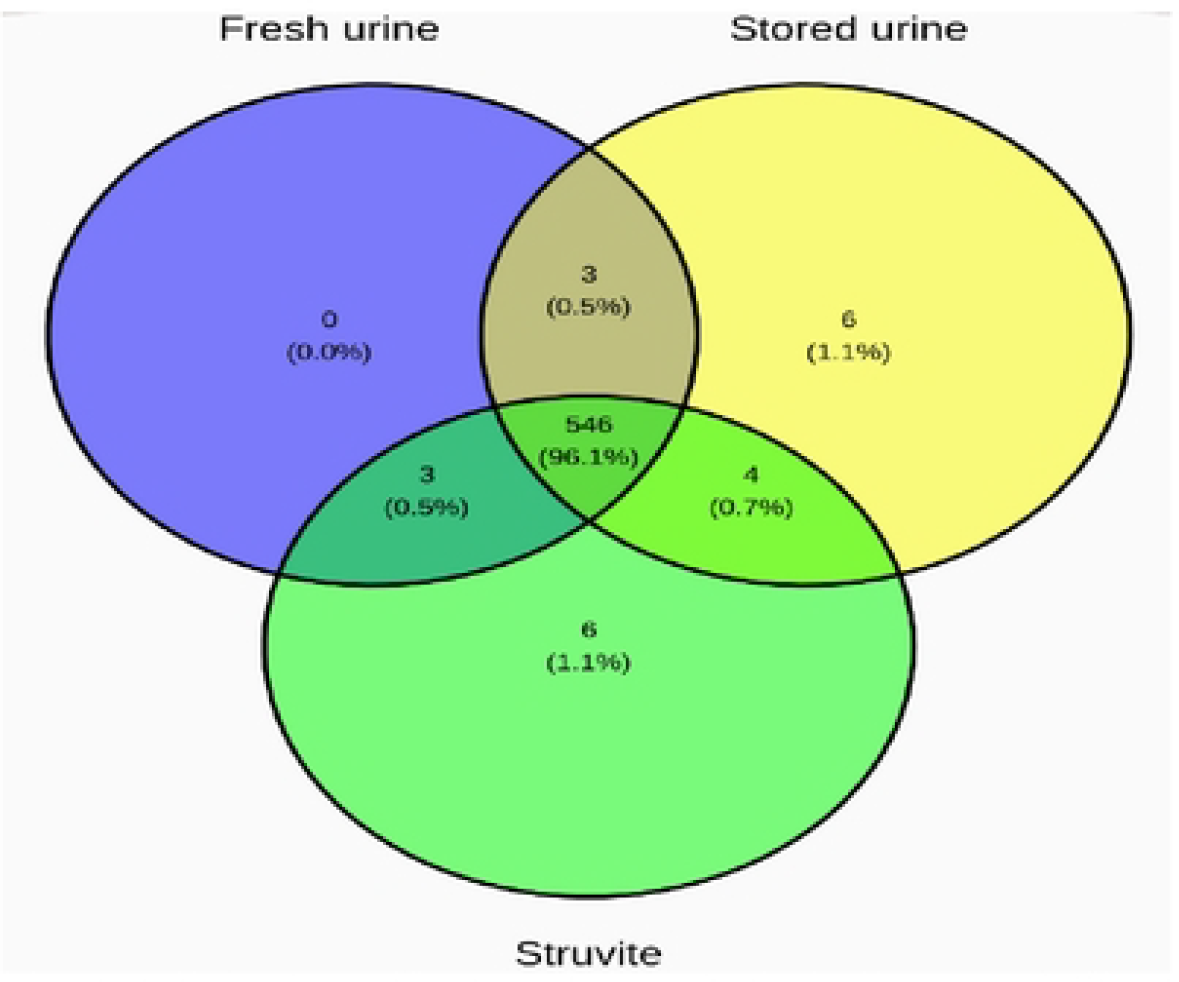
Count and percentages of shared genera in the three samples

### Notable antimicrobial resistance genes in the samples

The fresh urine sample harbored genes and mobile genetic elements responsible for resistance against gentamicin (*aac* family), streptomycin *(str)*, ESBL (*BLA-OXA)*, trimethoprim (*dfrA)*, erythromycin *(erm)* and macrolids (*erm)*. Diverse groups of resistance genes appeared in the struvite sample such as kanamycin (*ant(2’’)* family, *aph(3)1b*), ESBL (*BLA CARB* family), chloramphenicol (*cat* family), macrolide (*erm* family) and tetracycline (*tetD* and *H*) (Fig. 7). All of the detected resistance genes in the stored urine (*sul, cat, aph* and *aac* members) were not detected in the struvite sample while most of the genes from the fresh urine persisted in the struvite samples (Fig. 7).

**Figure 7.**
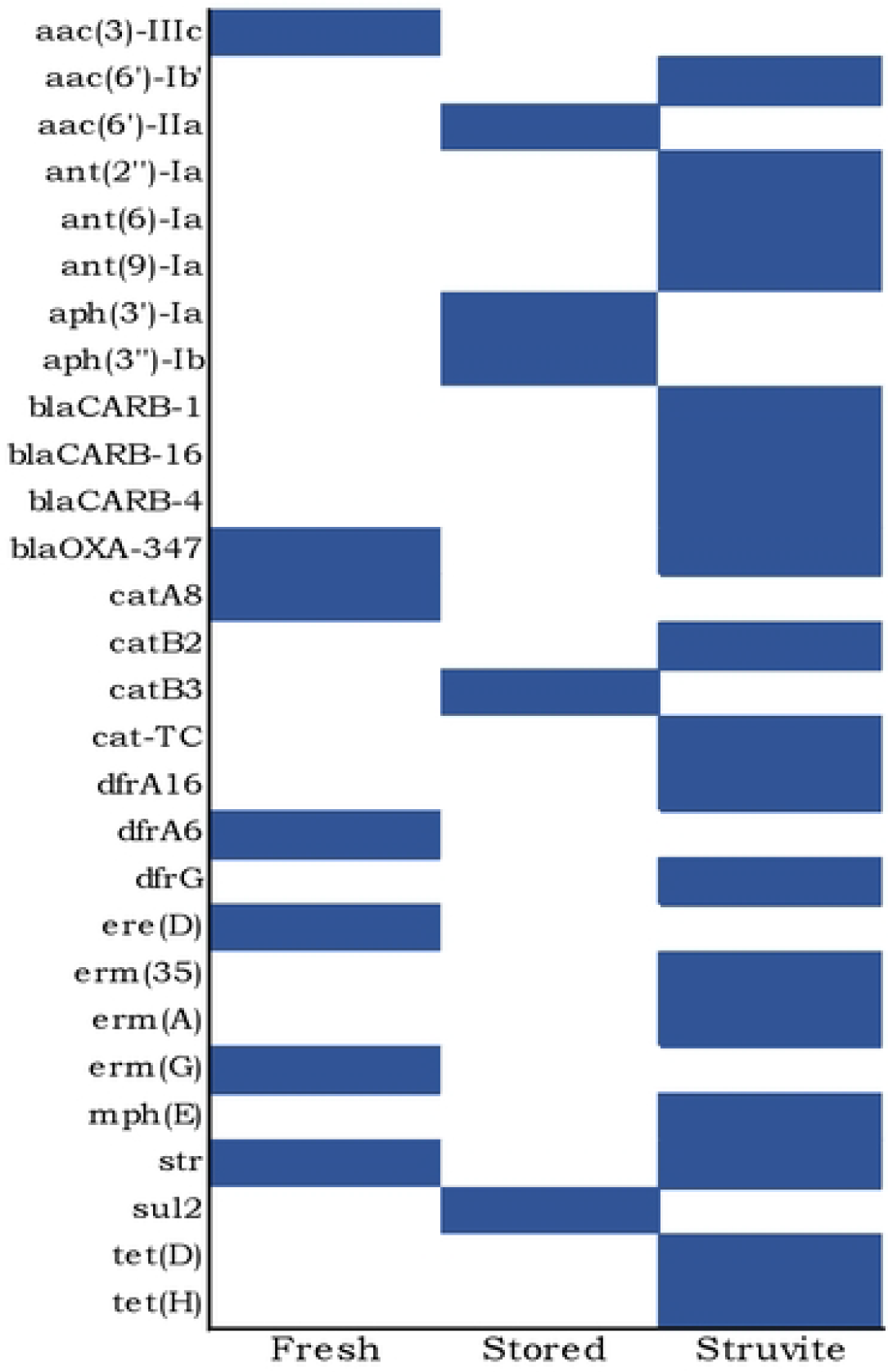
MGE-mediated antimicrobial resistance genes in the fresh, stored urine and struvite samples

### Diversity and coverage of microbial dynamics in the struvite production process

Shannon and Simpson diversity indices showed that the fresh urine diversity index is 2.8. The richness and abundance-based coverage estimator (ACE) of the fresh urine sample were 880 (ranging between 780 – 960) and 825 (810-830), respectively (Fig. 8). The stored urine had Shannon diversity index of 5.65 and ACE of 775 (745-780). The rarefaction curve indicated that in all the three samples the number of reads keep on increasing as the number of species increase, indicating that the sample has yet to cover the rare groups of the community (Fig. 9).

**Figure 8.**
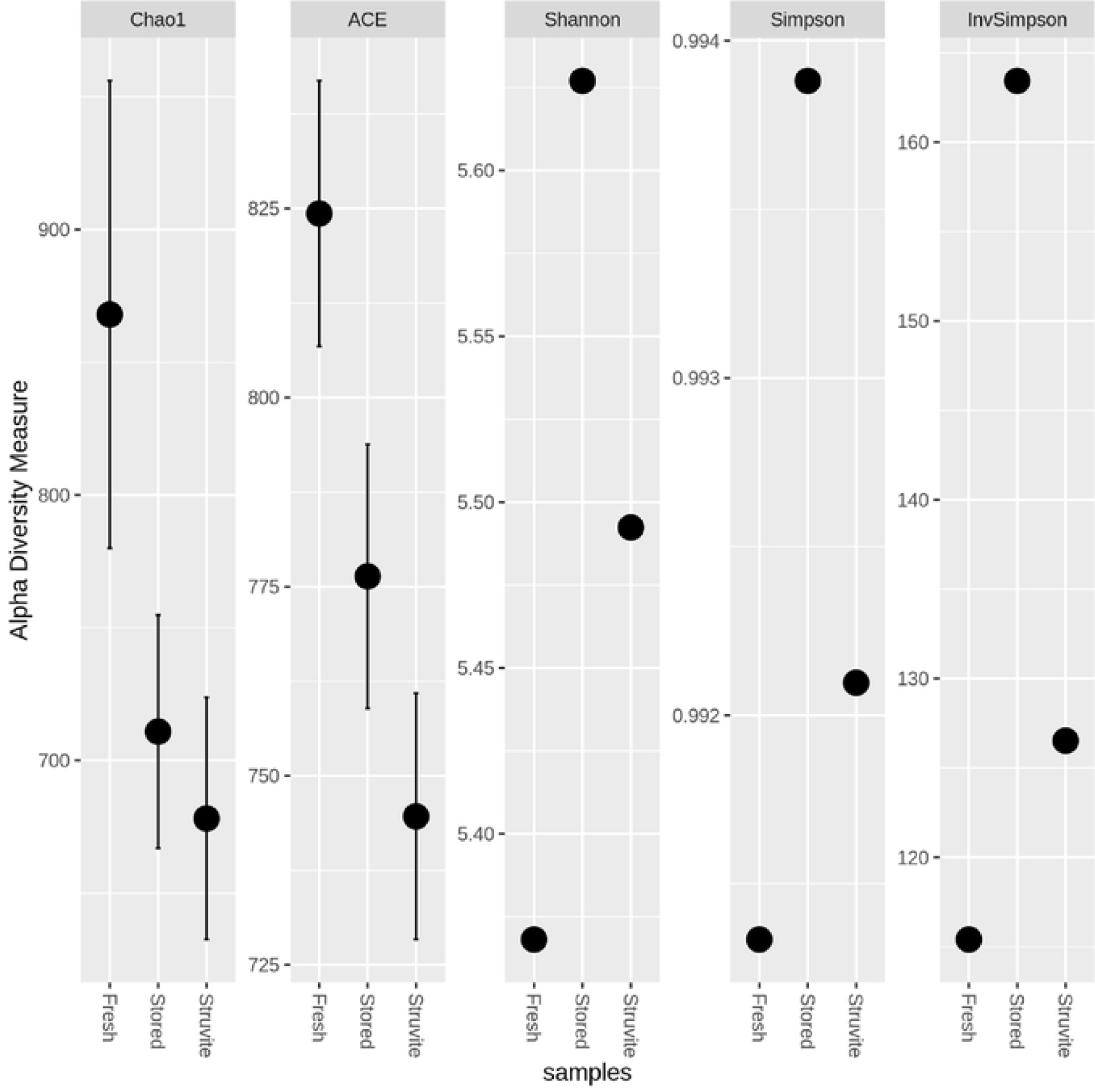
Diversity measures for microbes found in fresh urine, stored urine and struvite

**Fig. 9.**
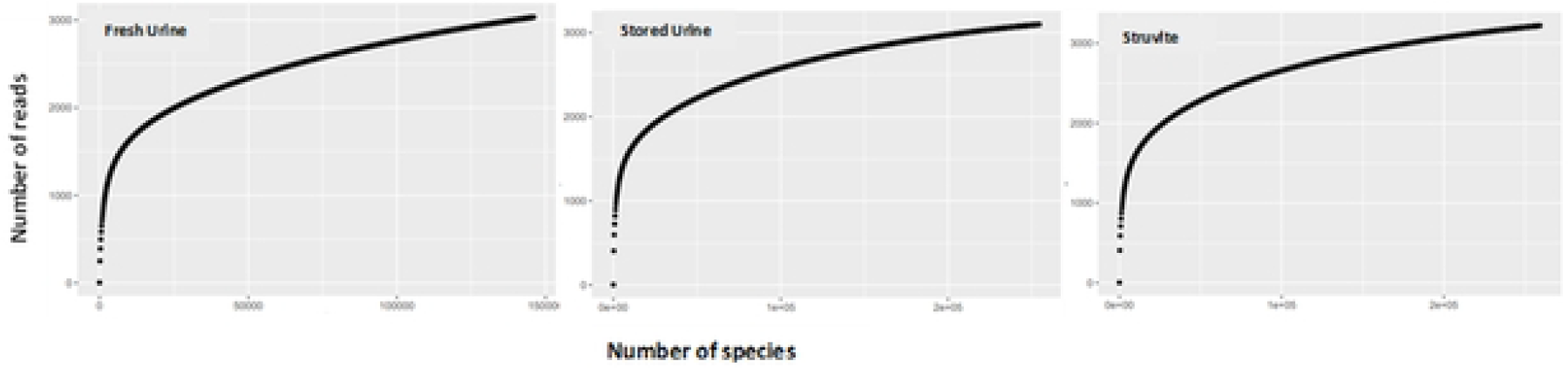
Rarefaction curve indicating the sample coverage and sequencing depth of the fresh, stored and struvite samples

## Discussion

The metagenomic reads generated from our samples had low proportion (< 0.5%) of unassigned reads indicating the power of the annotation analysis. Members of the phyla Pseudomonadota (formerly Proteobacteria), Bacillota (Firmicutes) and Actinomycetota were reported to dominate freshly collected urine using 16SrRNA-based urine microbiome study by (Nelson et al., 2010; Zuber et al., 2023). No phylum-level change and higher OTU-counts (not statistically significant) was observed in the stored urine than that of the fresh urine. As the stored urine has high pH (10) and ammonia level of nearly 4000mg/L (Lahr et al., 2016), change in the microbial composition and abundance is observed in the lower taxonomic levels. For example, notable decrease of members of alphaproteobacteria and gammaproteobacteria was observed in stored urine. Increase of the dominant members of bacteria in urine microbiota upon keeping for 4 days in room temperature was reported by (Jung et al., 2019), indicating that storage of urine even for a short period of time alters the microbial abundance.

The dominance of Bacillota (Firmicutes) and Bacteroidota in the struvite sample is indicative of the fact that struvite is not conducive for all bacteria except those with resistance mechanisms such as spore formation. Struvite is characterized by its low water activity compared to the liquid fresh or stored urine (Qiu et al., 2022), which makes it less conducive for microbial thriving except for those with survival mechanisms such as dormancy (Wetzel and McBride, 2020). In the study by (Lahr et al., 2016), the genera *Clostridia Tissierellia* and *Bacterodia* which belong to the dominant phyla in our study were reported to persist in the struvite samples. Further study is required on the possibility of regeneration to vegetative stage of the observed persistent genera in struvite.

The re-emergence of the genus *Pseudomonas* in the struvite sample after its marked decrease in the stored urine can be explained based on a recent work on the bacterial profile of kidney stones which revealed the presence of *Pseudomonas* in struvite stone (Halinski et al., 2023). The increase in abundance of *Shewanella* in the struvite sample can be attributed to the study by (Luo et al., 2018) which revealed the potential of *Shewanella oneidensis* strain as an inducer of struvite mineralization.

The storage time was higher in this study which in turn increase the exposure of the bacteria to higher amount of ammonia and alkaline pH, thus decreased the bacterial count (Goetsch et al., 2020; Lahr et al., 2016; Xu et al., 2022). The abundance of *Treponema* in the struvite sample may be speculated by the fatty acids present struvite may favor its presence (Salvatore, et al., 2013).

As the present study is metagenome-based, the fact that the majority of the bacteria genera being shared in all the samples is not indicative of the presence of live bacteria in all the samples. One of the limitations of metagenomic study is it provides circumstantial evidence on the live presence of the identified bacterial taxon.

The detection of diverse groups of Mobile genetic elements (MGE)-based resistance genes in the struvite sample is alarming which requires attention. Our previous report also indicated the abundance of phage- and plasmid-derived antimicrobial resistance genes (ARG) in the stored urine and struvite samples (Woldeyohannis and Desta, 2023). In this study, the MGE-based ARG involved insertion sequences in addition to plasmids. Explanation with the approach of ARG tracking is required to get a plausible explanation of such abundance in the struvite before moving forward with promoting the product for agricultural input.

Higher diversity and lower richness were observed in the stored urine sample than the rest of the samples and this is mainly attributed to the fact that urine as it comes out of the body is characterized by its low biomass (Jung et al., 2019). The stored urine exhibited the lowest evenness than the fresh urine and the struvite, which could be as a result of the dominant bacteria being perished by the alkaline pH and accumulation of ammonia during storage for an extended period of time (Lahr et al., 2016). This study revealed the optimized storage time (20 days) also contributed for the auto-sanitization of the urine from the majority of the bacteria.

In conclusion, optimal process parameters in the struvite production focuses only on the yield rather than safety. In our study, struvite produced with optimal process parameters may be precarious when it comes to safety and public health risks. We recommend culture-based study on the regeneration potential of members of the phyla Bacillota and Bacteroidota during application of struvite on soil. As the process of urine collection does not follow the clean-catch approach, we recommend complementary urine sanitization methods to be considered in struvite production. Furthermore, the drying process of the struvite should be given due consideration with regard to ARG minimization.

## Acknowledgement

The authors would like to thank Birtukan Getnet and Mr. Johanny Girma for urine collection and struvite processing. The financial support is from thematic research grant of Addis Ababa University.

